# Experimental rewilding of stickleback drives phenotypic shifts that oppose long-term evolutionary trajectories in Daphnia in Alaskan lakes

**DOI:** 10.1101/2025.10.15.682675

**Authors:** Stephanie M. Tran, Matthew R. Walsh

## Abstract

Trophic rewilding restores top-down interactions and may, in turn, drive novel patterns of evolutionary change. Predatory Northern pike invaded lakes in Alaska and eliminated native fish fauna. In 2018, pike were extirpated to ‘reset’ these ecosystems. Contrasting trophic ecomorphs (benthic vs. limnetic) of three-spine stickleback were then re-introduced as part of whole-lake rewilding experiments. We examined the top-down effects of stickleback on phenotypic change in *Daphnia* over three years in lakes that naturally contained sticklebacks and compared such trends with the experimental lakes. We observed consistent phenotypic differences between lakes with naturally occurring populations of benthic vs. limnetic stickleback. The patterns of phenotypic divergence indicate that predatory selection is stronger on *Daphnia* in lakes with benthic stickleback. The trends in the experimental lakes were temporally variable and opposed the trends observed in natural lakes. Rewilding provides insights into the inconsistencies that can manifest between short- and long-term influences of natural selection.

## INTRODUCTION

Rewilding is an ecosystem restoration strategy introduced in the late 20^th^ century that is designed to restore biodiversity and facilitate ecosystem stability (Svenning *et al*. 2016, 2024). Trophic rewilding approaches are aimed particularly at restoring top-down interactions. This approach alters the nature of exploitative and competitive interactions and has widespread cascading ecological consequences throughout the food web (Atkinson *et al*. 2024; Svenning *et al*. 2016). By altering top-down interactions, trophic rewilding therefore has the potential to modify selection and drive evolutionary changes at lower trophic levels. However, most work on rewilding has focused on large mammals and reptiles (tortoises) in terrestrial habitats (Svenning *et al*. 2016). Replicated rewilding experiments conducted in nature are rare but present the opportunity to address questions related the direction, tempo and repeatability of evolution.

*Esox lucius* (northern pike) are apex predators that have been introduced into lakes throughout the United States and have been linked to the reduction of native species in aquatic systems (Carim *et al*. 2022; Zelasko *et al*. 2016). Although indigenous north and west of the Alaskan Range, northern pike were introduced into numerous lakes on the Kenai Peninsula of Alaska beginning in the 1950’s. Since their introduction, northern pike extirpated or diminished populations of native fish such as *Oncorhynchus mykiss* (rainbow trout), *O. kisutch* (coho salmon) and *Gasterosteus aculeatus* (three-spined stickleback) (Alaska Department of Fish and Game 2007; Haught & Von Hippel 2011; K.J. Dunker *et al*. 2022). In Fall 2018, the Alaska Department of Fish and Game (ADFG) treated nine of these lakes with rotenone to cull and remove northern pike to allow for rewilding to restore native fish diversity.

In June 2019, >10,000 three-spined stickleback were reintroduced into the lakes that experienced the recent removal of pike to begin the process of whole ecosystem restoration (Dunker *et al*. 2020; Hendry *et al*. 2024). Three-spined stickleback have diversified into two phenotypic morphs with distinct trophic morphologies correlating to the characteristics of lakes they reside in. This includes: (i) ‘limnetic’ stickleback in large, deep lakes that exhibit traits specialized for foraging on pelagic zooplankton (i.e., higher jaw protrusion, numerous, fine gill rakers, larger eye size, and shallower heads and bodies), and (ii) ‘benthic’ stickleback in small, shallow lakes which exhibit traits specialized for foraging on macrobenthos in shallow water (e.g., fewer, widely spaced gill rakers, tougher pharyngeal teeth, and deeper heads and bodies) (Des Roches *et al*. 2013; McGee *et al*. 2013; Willacker *et al*. 2010). Divergent morphological forms of these stickleback were split and distributed evenly amongst the eight lakes. Four lakes received limnetic stickleback, four lakes received benthic stickleback, while a ninth lake received both limnetic and benthic stickleback [though a second lake ultimately received benthic and limnetic stickleback – see (Hendry *et al*. 2024)]. The two morphs of this species have been shown to differentially affect zooplankton community structure and processes through trophic interactions (Des Roches *et al*. 2013; Harmon *et al*. 2009). As a result, the presence of these divergent forms of stickleback that forage in contrasting environments may alter selection in a divergent manner for their prey.

Here we tested if the experimental rewilding of stickleback drives evolution in their prey. After the rotenone treatment, *Daphnia* naturally recolonized the rewilded lakes via resting eggs found in lake sediment. *Daphnia* (waterfleas) are model organisms for eco-evo research due to their pivotal role as keystone species and their ability to undergo rapid evolutionary change (Ebert 2022; Reynolds 2011). As one of the dominant zooplankton grazers of phytoplankton, they exhibit top-down effects on phytoplankton biomass, nutrient cycling, and primary production (Bergquist & Carpenter 1986; Brett *et al*. 1994; Wagner *et al*. 2013). At the same time, *Daphnia* are heavily predated by invertebrates and planktivorous fish, including the three-spined stickleback (Hambright & Hall 1992; Visser 1982). Much research has shown that differences in predation can drive shifts in *Daphnia* behavior, life history and morphology (Walsh & Post 2011, 2012; Landy *et al*. 2020). For example, having longer tail spines are useful by prolonging handling time, which decreases predator foraging efficiency and provides greater opportunity for prey to escape (Black & Dodson 1990; Caramujo & Boavida 2000).

Our study had two goals: First, we tested for phenotypic divergence between populations of *Daphnia* from lakes in Alaska with naturally occurring populations of limnetic and benthic stickleback. Such comparisons will provide information on the relative endpoints of the evolutionary process. Secondly, we compared these patterns to lakes with the rewilded populations of stickleback. We assessed the traits of *Daphnia* over three consecutive years (2022-24) starting three years after the initiation of the rewilding experiments. The results stemming from the rewilded lakes allow us to determine the repeatability and consistency when *Daphnia* initially adapt to the presence of a novel predator. Because limnetic sticklebacks are open-water specialists that forage primarily on zooplankton, we expected that limnetic sticklebacks should lead to increased mortality and/or stronger selection on *Daphnia* (i.e., smaller body size, bigger eye size, longer tail-spine, and a stronger anti-predator behavior). However, the presence of limnetic vs. benthic stickleback also covaries with additional factors that may influence selection on *Daphnia*. Particularly, lakes with benthic stickleback are typically shallower with less deep-water refuge for *Daphnia*. Furthermore, the rewilded lakes vary along a size gradient (Hendry *et al*. 2024) that may further influence the trajectory of evolution.

## MATERIALS AND METHODS

### (a) Study Lakes and Sampling

These study lakes are a part of a larger eco-evo lake restoration experiment on the Kenai Peninsula of Alaska. The details of the methods behind setting up these rewilded lakes are fully described in (Hendry *et al*. 2024). We studied evolutionary divergence in *Daphnia* from seven of these lakes in Alaska during three sampling periods in July 2022, 2023, and 2024. We included five ‘experimental’ or rewilded lakes that experienced the reintroduction of stickleback. We used a subset of the full set of rewilded lakes because these are the lakes that sustained populations of *Daphnia. Daphnia* were absent from the other rewilded lakes. Lakes that experienced the reintroduction of stickleback include two limnetic lakes (Hope, Crystal), two benthic lakes (CC, Leisure), and one lake with both populations of stickleback (Loon). Our study also included two ‘natural’ lakes that were never invaded by pike and contained long-standing populations of stickleback and *Daphnia*. The natural lakes included one lake with limnetic stickleback (Spirit) and one lake with benthic stickleback (Watson). All seven lakes were sampled in 2022-24, although *Daphnia* were absent at the time of sampling in Hope Lake in 2023 and Crystal Lake in 2024. It is also important to note that there are currently different species of *Daphnia* in the natural vs. rewilded lakes. The natural lakes contain *D. mendotae*, while *D. longiremis* is present in the rewilded lakes.

Zooplankton samples were collected using a plankton net from the greatest depth of each lake. Samples were then immediately transferred to a nearby residence to isolate *Daphnia*. From each sample, we took images to quantify *Daphnia* body size, eye size, tail-spine length and performed experiments to assess anti-predator behavior (phototactic responses). All behavioral assays and images (see below) were completed within 1 to 5 hours of collection for each population. The data are uploaded on Dryad (Tran and Walsh 2025).

### (b) Diel Vertical Migration (DVM) Trials

*Daphnia* naturally exhibit an anti-predator response where they migrate to the bottom of lakes during the day to avoid visual fish predators (Cousyn *et al*. 2001; Forward 1988). This anti-predator behavior is assessed by measuring phototactic behavior via vertical migration trials in experimental columns. 9 to 14 individuals (overall average of 10.0 ± 0.3 individuals per trial) were randomly selected from lake samples and acclimated in a 38-cm vertical glass column enclosed in a darkened station for 7 to 14 minutes (overall average of 9.5 ± 1.1 minutes per acclimation). The columns were enclosed in a rectangular PVC structure (30cmx30cmx50cm) that was wrapped in black plastic to ensure that the acclimating columns were in complete darkness. The column was divided into three sections: a 3-cm lower section, a 10-cm middle section, and a 12-cm upper section. After the acclimation period, a diffuse light above the column was switched on (Fenix LD15R) and the number of individuals in each section was recorded every minute for 7 minutes. The light entering each enclosure was quantified each day prior to conducting trials to ensure consistency in light for all trials (Light meter:). For each combination of lake and year, 14 to 29 trials were run (overall average of 17 ± 8 trials per lake, total no. of trials =372). We calculated the phototactic index (PI) for each minute, then calculated the average PI over the course of the 7-minute trial. The same set-up and light sources were used for all trials for all three years of data collection.

The phototactic index is defined as, PI = (U-L)/(U+M+L), where U, M, and L are the number of individuals in the upper, middle, and lower sections of the column. Values range from -1 (negatively phototactic) to +1 (positively phototactic). Each trial produced one estimate of PI.

### (c) Morphological Traits

To quantify morphological anti-predator defenses, we imaged 19 to 52 individuals per sample (32 ± 13 individuals per lake per year, total n=674 images) using a Leica EZ4 W microscope with integrated 5.0 Mega Pixel CMOS WiFi camera. Image J software was used to measure body size, eye size, and tail-spine length. Body and eye size were measured as the longest distance between any two points using Feret’s Diameter and tail-spine length was measured from the base at the body to the end of the tail.

### (d) Statistical Analyses

We tested for trait differences in *Daphnia* from lakes with benthic and limnetic sticklebacks using ANOVA (Type III). We analyzed the data from the natural and rewilded lakes separately because the natural and rewilded lakes contain different species of *Daphnia*. Plus, *Daphnia* in the natural vs. rewilded lakes have experienced contrasting durations of predation by stickleback. All statistical analyses were performed using R Studio Version 2023.12.1+402 (Posit team 2024).

First, we tested for differences in morphology. We natural ln-transformed body size, eye size, and tail-spine length to better meet assumptions of normality and homogeneity of variances. We included stickleback ecotype (benthic, limnetic), year of sampling (2022, 2023, 2024) and the ‘ecotype by year’ interaction as our fixed effects. We included body size, absolute eye size, and absolute tail-spine length as our dependent variables. We then tested for ‘relative’ eye size and tail-spine length by including body size as a covariate.

We also tested for differences in phototactic behavior. We used the overall average phototactic index (PI) per trial as our dependent variable. The stickleback ecotype, year of sampling, and the ‘ecotype by year’ interaction were included as fixed effects. The number of individuals per trial was included as a covariate in the analyses of phototactic behavior. Note: phototactic behavior was only analyzed for years 2022 and 2023 in the rewilded lakes due to low abundances or absence of *Daphnia* from lakes with limnetic stickleback in 2024. *Daphnia* was absent from Crystal Lake and observed at low abundances in Hope Lake in 2024. This allowed us to assess the morphological traits of Daphnia in Hope Lake but the low sample size precluded assessment of PI.

## RESULTS

### Natural lakes

We observed significant (p<0.05) differences in body size, absolute and relative eye size, absolute and relative tail-spine length, and phototactic behavior between ecotypes (Table 1). On average across years, *Daphnia* from the lake with benthic stickleback were 35% more negatively phototactic than *Daphnia* from the lake with limnetic stickleback. *Daphnia* from the lake with benthic stickleback were 8% smaller than the *Daphnia* from the lake with limnetic stickleback lake but exhibited a relative eye size and tail-spine length that were 6% and 11% larger, respectively (Fig. 1). The differences in body size and absolute eye size varied across years as the ecotype x year interaction was significant (Table 1). *Daphnia* from the lake with benthic stickleback were smaller in size when compared with *Daphnia* from the lake with limnetic stickleback in 2022 and 2023. Such size differences increased in 2024. There were no differences in absolute eye size in 2022 and 2023, but *Daphnia* eye size was smaller in the lake with benthic stickleback in 2024. The ecotype x year interactions were not significant (p>0.05) for absolute and relative tail-spine length, relative eye size, and phototactic index (Table 1).

**Figure 1.**
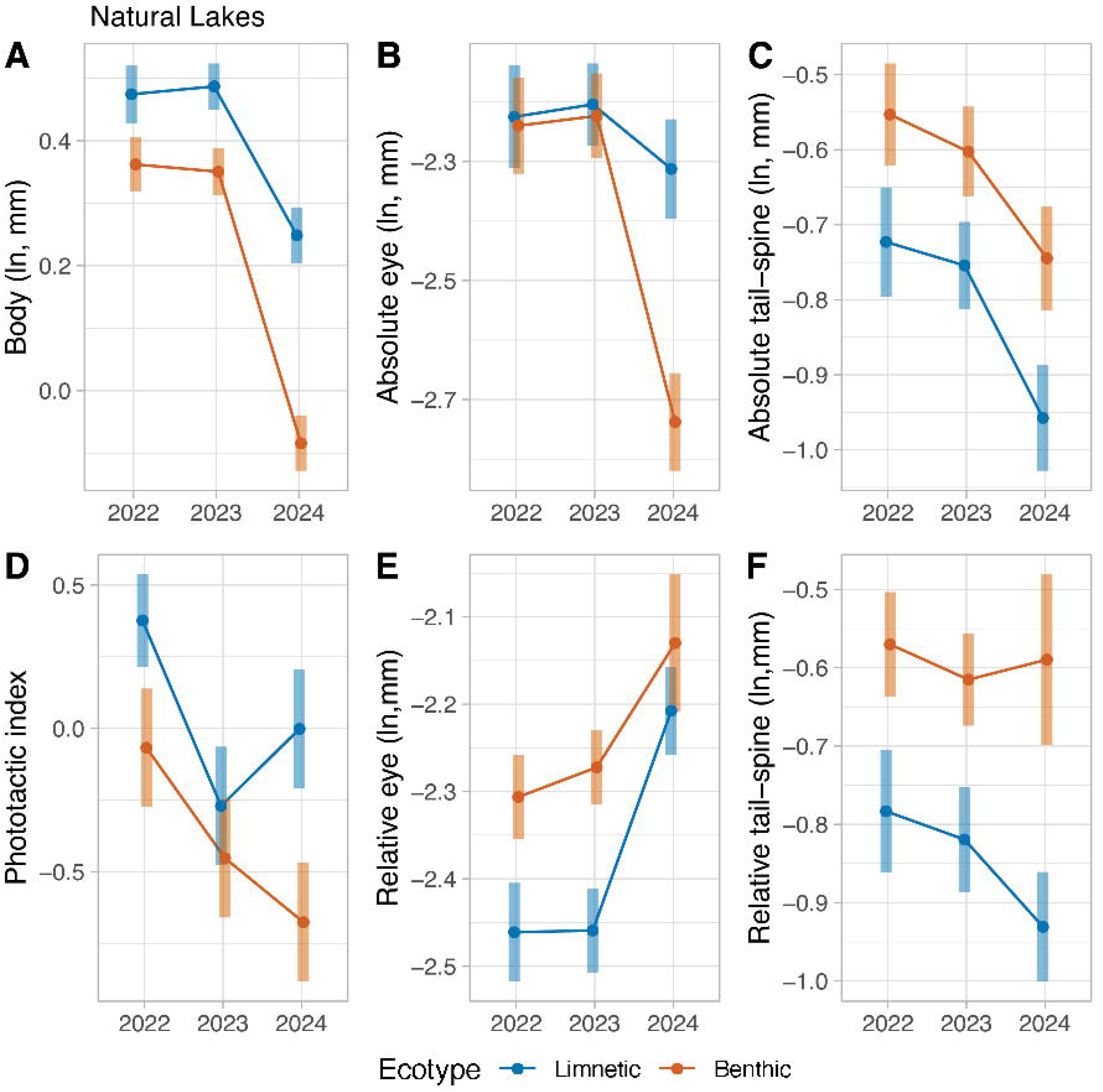
Estimated marginal means (EMMs) in *Daphnia* body size, eye size, tail-spine length, and phototactic behavior in natural lakes across the years. (A-F) Natural lakes. Body size, absolute eye and tail-spine lengths are in ln-mm. Phototactic index is the average PI. Note that there is only one lake that contains both benthic and limnetic stickleback in the natural lakes. Error bars denote confidence interval.

### Rewilded lakes

The differences in trait values between lakes with contrasting populations of sticklebacks varied across years. This is because we observed a significant (p<0.05) ecotype x year interaction for body size, absolute and relative eye size, absolute (but not relative) tail-spine length, and phototactic index (Table 2). Similar to the trends observed in natural lakes, *Daphnia* from lakes with benthic stickleback were initially (in 2022) smaller, with a smaller absolute eye size, and were more negatively phototactic when compared with *Daphnia* from lakes with limnetic stickleback (Fig. 2). But such differences disappeared (in 2023 for PI) or were reversed (body and eye size in 2023-24). The differences in relative eye size were more consistent. Overall, *Daphnia* from lakes with benthic stickleback exhibited a relative eye size that was 6% and 7% larger than *Daphnia* that experienced the introduction of limnetic stickleback or both benthic and limnetic stickleback (Loon Lake), respectively. The ecotype x year interaction was significant for this trait because the differences in relative eye size varied between the lakes with limnetic stickleback and both ecomorphs across years (Fig. 2).

**Figure 2.**
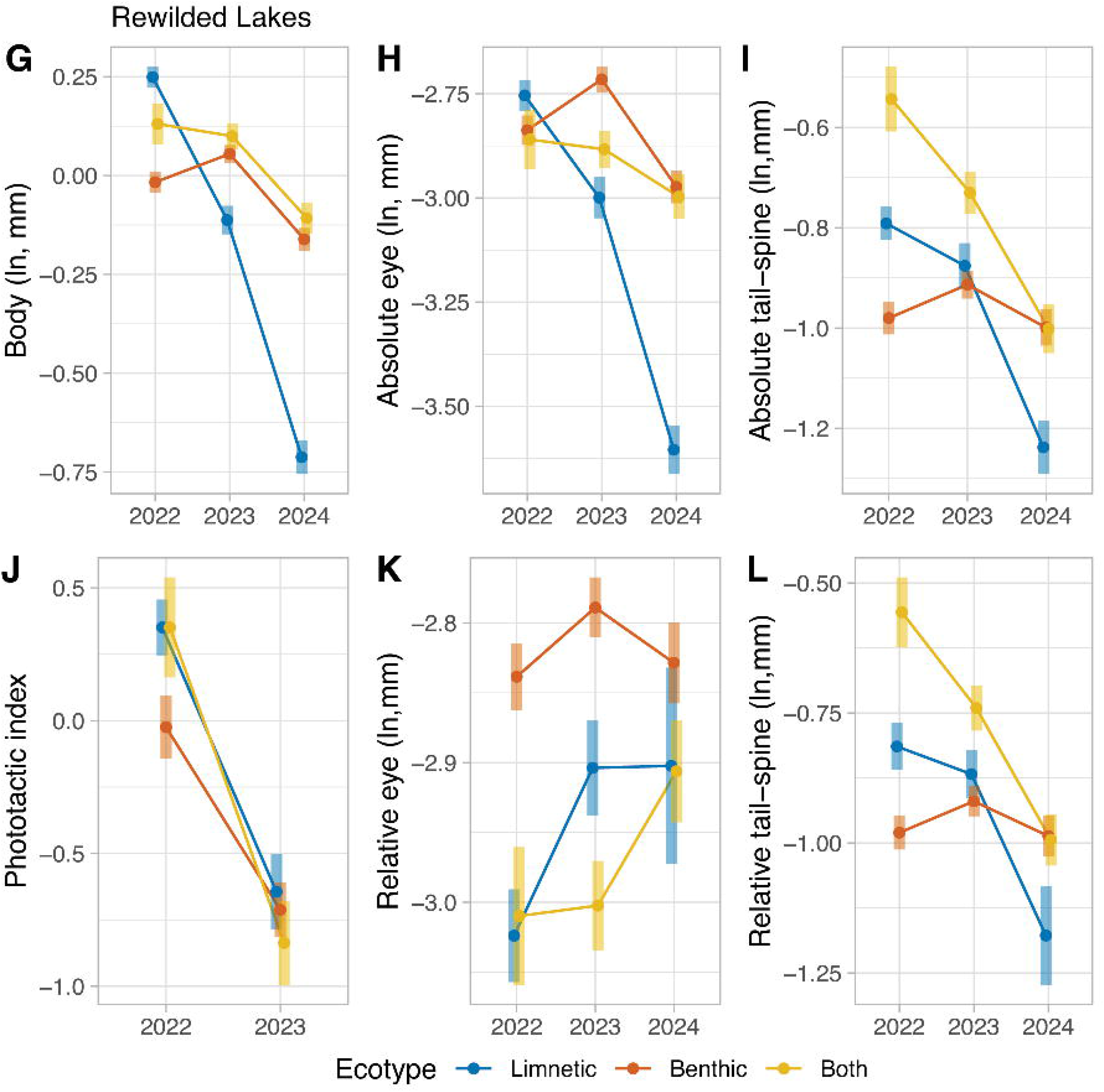
Estimated marginal means (EMMs) in *Daphnia* body size, eye size, tail-spine length, and phototactic behavior in experimental (rewilded) lakes across the years. (G-L) Natural lakes. Body size, absolute eye and tail-spine lengths are in ln-mm. Phototactic index is the average PI. In 2023 and 2024, *Daphnia* was observed in one lake containing rewilded limnetic stickleback. In 2024, phototactic behavior was not collected for limnetic rewilded lakes. Error bars denote confidence interval.

## DISCUSSION

We observed a temporally consistent pattern of phenotypic divergence between lakes with naturally occurring populations of benthic and limnetic stickleback. *Daphnia* from a lake with benthic stickleback were more negatively phototactic, had a smaller body size, relatively larger eyes, and longer tail spines than a lake with limnetic stickleback. In the rewilded lakes, the phenotypic differences between lakes with introduced benthic vs. limnetic stickleback paralleled the natural lakes in 2022 (except for tail-spine length). After 2022, such differences largely disappeared or were reversed except for relative eye size. That is, the patterns of divergence in the rewilded lakes were temporally variable. These results beg three questions: (1) Why are benthic stickleback associated with a phenotypic trajectory that is opposite to our *a priori* expectations? (2) Why did we only see consistent shifts in rewilded and natural lakes in relative eye size? and (3) What explains the volatile trends observed in the rewilded lakes?

In prior studies, there has been strong support for the idea that increased predation selects for smaller body size, relatively larger eyes, and longer tail spines in *Daphnia* (Beston *et al*. 2019; Beston & Walsh 2019; Nagano *et al*. 2023; Spaak & Boersma 1997). *Daphnia* in the natural lake with benthic stickleback were more negatively phototactic, smaller, and had relatively larger eyes and longer tail-spines (Fig. 1). These results indicate that benthic stickleback impose stronger predatory selection on *Daphnia* in the long-standing natural lakes included in our study. It is important to note that benthic and limnetic stickleback covary with other ecological and bathymetric features of lakes that could modify the nature of selection on the traits of *Daphnia*. Benthic sticklebacks are typically found in lakes that are shallower compared to lakes with limnetic stickleback. This is true for the lakes included in this study as the maximum depth of Watson Lake is 4 meters, compared to the maximum depth of 20 meters in Spirit Lake. In shallow lakes that lack vertical habitat partitioning, *Daphnia* may have more limited ability to avoid intense bouts of predation by both planktivorous fish (i.e., stickleback) and benthic predatory invertebrates such as *Chaoborus* and *Leptodora*. A potential alternative explanation, at least for body size, is intraspecific competition. A smaller body size, as seen in *Daphnia* from Watson Lake, may also be due to nutrient limitations coupled with increased intraspecific competition (Rautio *et al*. 2011; Rautio & F. Vincent 2007; Rautio & Vincent 2006). Benthic lakes have an abundance of benthic mats due to nutrient-rich sediment and high benthic productivity (Ask *et al*. 2009; Cazzanelli *et al*. 2012; Mariash *et al*. 2011, 2014; Quesada *et al*. 2008; Rautio *et al*. 2011; Rivera Vasconcelos *et al*. 2018). Zooplankton have documented to feed on lower quality, benthic organic matter in lieu of phytoplankton in such circumstances (Cazzanelli *et al*. 2012; Mariash *et al*. 2011). It seems likely that trends in the natural lake with benthic stickleback are a byproduct of benthic stickleback being in small, shallow lakes that reduce deep water refuge and the ability of *Daphnia* to avoid being eaten. One caveat is that our natural lakes are not replicated at the ecotype level. Thus, the consistency of these trends across multiple lakes with benthic vs. limnetic stickleback is unknown at this time.

The trend of having relatively larger eyes in rewilded lakes with benthic stickleback mirrors the patterns observed in *Daphnia* in natural lakes. This suggests that there are fitness advantages to having larger eyes in lakes with benthic sticklebacks. In particular, larger eyes may aid in avoiding predation. To avoid lethal attacks, prey must first rely on sensory systems to successfully detect predators while performing other tasks. Sensory systems collect internal and external information and then mediate behavioral, physiological, and morphological changes in an organism. It is known that environmental visual cues such as light availability (Hall 2008; Lisney & Collin 2007; Thomas *et al*. 2002), predation (Lönnstedt *et al*. 2013; Svanbäck & Johansson 2019; Vinterstare *et al*. 2020), and foraging (Merry *et al*. 2011) affect eye evolution – particularly eye size. *Daphnia* have a compound eye that is capable of optomotor responses to assess these cues (Consi *et al*. 1990; Frost 1975). Eye size in *Daphnia* is highly plastic and has been shown to vary in response to low-quality food (Brandon & Dudycha 2014; Walsh & Gillis 2021), light (Howell *et al*. 2023), and has major implications for fitness (Beston *et al*. 2019; Beston & Walsh 2019; Brandon *et al*. 2015; Kessler & Lampert 2004). Investment in sensory systems (i.e., eye tissue) is energetically expensive (Niven 2015; Niven & Laughlin 2008), potentially incurring tradeoffs with other antipredator defenses due to constraints on resource availability (Kikuchi *et al*. 2023). Due to the link between visual systems, fitness, and enhanced survival, eye size appears to be a driving factor in adapting to lakes with sticklebacks, especially with benthic stickleback. Our results suggest that increased predator detection is important in the early stages of selection following the rewilding of stickleback.

The trait differences observed in the natural lakes presumably represent the end point of the evolutionary interactions between stickleback and *Daphnia* (assuming relatively consistent year to year environmental conditions). The trends in the rewilded lakes were less consistent (Fig. 2). Furthermore, the introduction of stickleback was also associated with rapid shifts in species composition. Prior to our 2022 sampling, the rewilded lakes were dominated by large-bodied zooplankton characteristic of lakes without planktivorous fish such as *D. middendorffiana* and *Hesperodiaptomus septentrionalis* (personal obs.). The large *Daphnia* (*D. middendorffiana)* were rapidly replaced by a much smaller species of *Daphnia* (*D. longiremis)* by 2022. This initial trend towards small-bodied zooplankton following the sudden appearance of planktivorous fish has been documented due to initial high levels of planktivory (Hyatt *et al*. 2021; Mittelbach *et al*. 1995, 2006). Furthermore, the initial patterns of phenotypic divergence observed in 2022 provided evidence that predatory selection was stronger in the rewilded lakes with benthic stickleback (i.e., *Daphnia* were smaller and more negatively phototactic). But such trends were largely reversed over the past three years. Notably, we saw a sharp decrease in body size for each limnetic rewilded lake and a decline in abundance over the years. We did not observe *Daphnia* in Hope Lake in 2023 but were able to sample a scarce quantity of smaller individuals in 2024. Similarly, after a decrease in body size, we did not collect any *Daphnia* in Crystal Lake in 2024. These trends and declines in abundances are not simply an effect of lake size. Hope and Crystal Lake are larger and deeper than one of the lakes with benthic stickleback (CC). More importantly, these trends provide evidence that there has been a temporal shift in the strength of selection; predatory selection appears to now be stronger in the lakes with limnetic. The question now is whether these trends will reverse over time to match the consistent patterns documented in the natural lakes.

This study highlights how intraspecific variation in planktivorous predators can drive phenotypic divergence in their prey. Interestingly, there is a disconnect in the patterns of phenotypic divergence in lakes with longstanding populations of benthic and limnetic stickleback versus lakes that experienced the recent rewilding of divergent stickleback morphs. It remains to be seen if the patterns of phenotypic change between the natural and rewilded lakes will ultimately converge or remain fundamentally distinct. These rewilding experiments provide further insight into how initial trajectories of evolution could potentially be opposite to the ultimate trajectory of change.

## Acknowledgements

We thank Meghan Korte, Caleb Miller, Kevin Tran, Yasmine Castillo, Amanda Stoner, and Marcus Lee for help in the field. The Alaska Department of Fish and Game granted collection permits. The University of Texas at Arlington provided funding support.

## TABLE AND FIGURE CAPTIONS

Table 1. Result of analyses for body size, absolute and relative eye size, absolute and relative tail-spine length (ln, mm), and phototactic index (PI) for 2022-24 in the natural lakes. Note: There only is one lake for each ecotype (Spirit, Benthic). Significant values (p<0.005) are bolded.

Table 2. Result of analyses for body size, absolute and relative eye size, absolute and relative tail-spine length (ln, mm), and phototactic index (PI) for 2022-23 in the rewilded lakes. Note: phototactic behavior was only analyzed for years 2022 and 2023 in the experimental lakes due to the absence of *Daphnia* from lakes with limnetic stickleback in 2024. Significant values (p<0.05) are bolded.

**Supplemental Figure 1**. Compiled, means of raw data for Daphnia body size, eye size, tail-spine length, and relative eye/tail residuals (mm) and phototactic index in natural and rewilded lakes by year (M-R). Open circles with dotted lines indicated natural lakes, closed circles with solid lines indicate rewilded lakes, and colors represent ecotype. Error bars denote confidence interval.

