## Supplemental Table 1 for "Experimental rewilding of stickleback drives phenotypic shifts that oppose long-term evolutionary trajectories in Daphnia in Alaskan lakes"

**Table 1:** Result of analyses for body size, absolute and relative eye size, absolute and relative tail-spine length (ln, mm), and phototactic index (PI) for 2022-24 in the natural lakes. Note: There only is one lake for each ecotype (Spirit, Benthic). Significant values (p<0.005) are bolded.

| **Natural lakes** | | **Body** | | | | **Absolute eye** | | | | | **Absolute tail-spine** | | | |
| --- | --- | --- | --- | --- | --- | --- | --- | --- | --- | --- | --- | --- | --- | --- |
|  |  | df | Denom.df | F | P-value | df | Denom. df | F | P-value | df | | Denom. df | F | P-value |
| Fixed effects | **Ecotype** | 1 | 200 | 122.98 | **<0.001** | 1 | 200 | 21.93 | **<0.001** | 1 | | 200 | 41.72 | **<0.001** |
|  | **Year** | 2 | 200 | 158.42 | **<0.001** | 2 | 200 | 37.02 | **<0.001** | 2 | | 200 | 20.95 | **<0.001** |
|  | **Ecotype × Year** | 2 | 200 | 15.09 | **<0.001** | 2 | 200 | 16.63 | **<0.001** | 2 | | 200 | 0.44 | 0.642 |
|  | | **Phototactic Index (PI) 22-24** | | | | **Relative eye** | | | | | **Relative tail-spine** | | | |
|  |  | df | Denomdf | F | P-value | df | Denom. df | F | P-value | df | | Denom. df | F | P-value |
| Covariates | **Body** |  |  |  |  | 1 | 199 | 372.85 | **<0.001** | 1 | | 199 | 12.65 | **<0.001** |
|  | **Individual** | 1 | 113 | 0.27 | 0.604 |  |  |  |  |  | |  |  |  |
| Fixed effects | **Ecotype** | 1 | 113 | 63.61 | **<0.001** | 1 | 199 | 32.31 | **<0.001** | 1 | | 199 | 55.10 | **<0.001** |
|  | **Year** | 2 | 113 | 36.91 | **<0.001** | 2 | 199 | 18.80 | **<0.001** | 2 | | 199 | 1.63 | 0.199 |
|  | **Ecotype × Year** | 2 | 113 | 7.135 | **0.001** | 2 | 199 | 2.55 | 0.081 | 2 | | 199 | 2.32 | 0.101 |
