## Supplemental Table 2 for "Experimental rewilding of stickleback drives phenotypic shifts that oppose long-term evolutionary trajectories in Daphnia in Alaskan lakes"

**Table 2.** Result of analyses for body size, absolute and relative eye size, absolute and relative tail-spine length (ln, mm), and phototactic index (PI) for 2022-23 in the rewilded lakes. Note: phototactic behavior was only analyzed for years 2022 and 2023 in the experimental lakes due to the absence of *Daphnia* from lakes with limnetic stickleback in 2024. Significant values (p<0.05) are bolded.

| **Rewilded lakes** | | **Body** | | | | **Absolute eye** | | | | **Absolute tail-spine** | | | |
| --- | --- | --- | --- | --- | --- | --- | --- | --- | --- | --- | --- | --- | --- |
|  |  | df | Denom. df | F | P-value | df | Denom. df | F | P-value | df | Denom. df | F | P-value |
| Fixed effects | **Ecotype** | 2 | 470 | 115.63 | **<0.001** | 2 | 470 | 123.65 | **<0.001** | 2 | 470 | 72.85 | **<0.001** |
|  | **Year** | 2 | 470 | 474.15 | **<0.001** | 2 | 470 | 194.44 | **<0.001** | 2 | 470 | 144.92 | **<0.001** |
|  | **Ecotype × Year** | 4 | 470 | 179.17 | **<0.001** | 4 | 470 | 80.08 | **<0.001** | 4 | 470 | 37.74 | **<0.001** |
|  | | **Phototactic Index (PI) 2022-23** | | | | **Relative eye:body** | | | | **Relative tail-spine** | | | |
|  |  | Df | Denom df | F | P-value | df | Denom. df | F | P-value | df | Denom. df | F | P-value |
| Covariates | **Body** |  |  |  |  | 1 | 469 | 562.12 | **<0.001** | 1 | 469 | 2.24 | 0.135 |
|  | **Individuals** | 1 | 183 | 0.99 | 0.321 |  |  |  |  |  |  |  |  |
| Fixed effects | **Ecotype** | 2 | 183 | 6.93 | **0.001** | 2 | 469 | 92.20 | **<0.001** | 2 | 469 | 57.45 | **<0.001** |
|  | **Year** | 1 | 183 | 276.57 | **<0.001** | 2 | 469 | 9.69 | **<0.001** | 2 | 469 | 39.94 | **<0.001** |
|  | **Ecotype × Year** | 2 | 183 | 6.86 | **0.001** | 4 | 469 | 7.33 | **<0.001** | 4 | 469 | 23.44 | **<0.001** |
