## Supplementary figures and images for "Experimental rewilding of stickleback drives phenotypic shifts that oppose long-term evolutionary trajectories in Daphnia in Alaskan lakes"

### Supplemental Figure 1

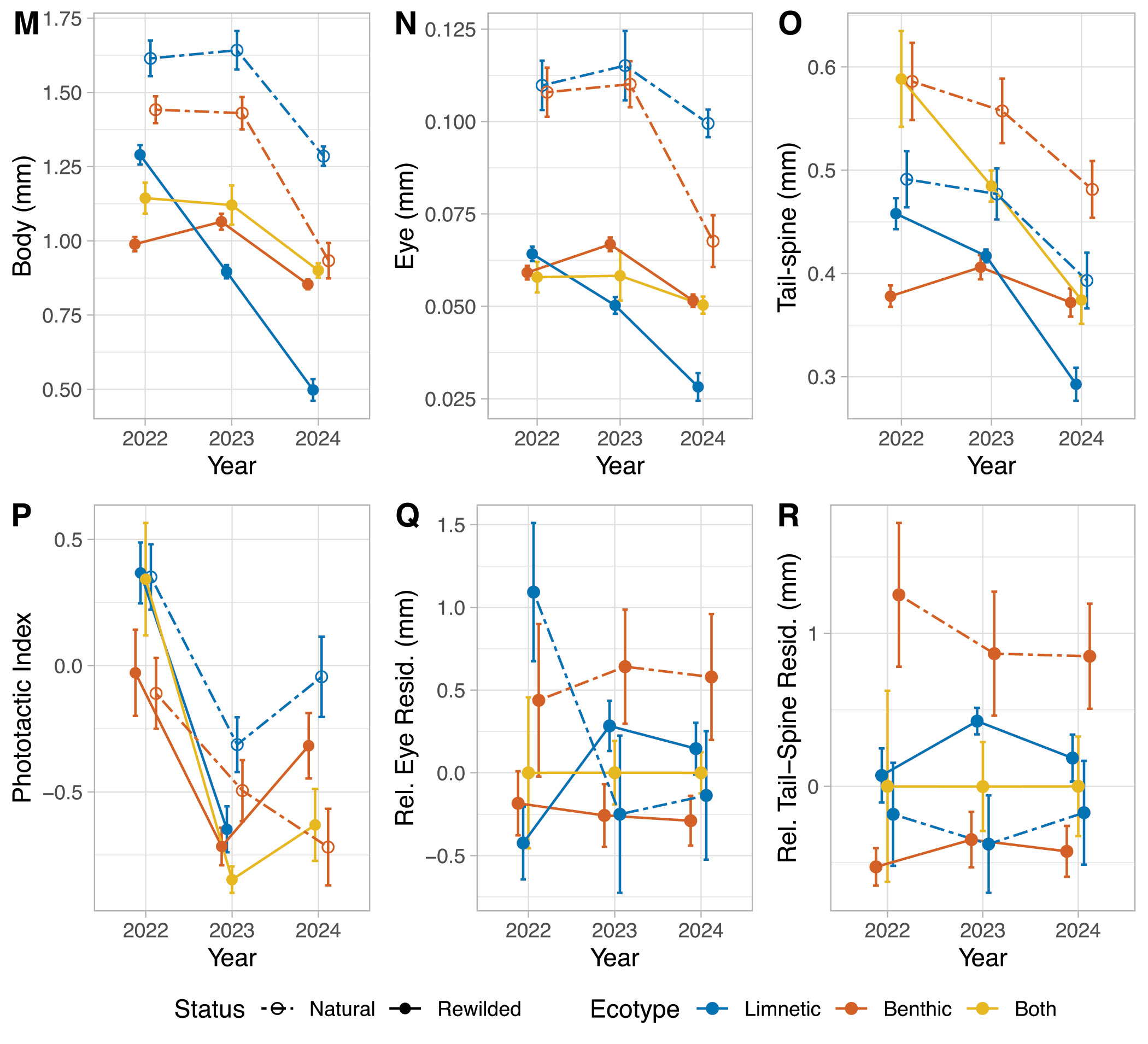
